# RevPert: predicting candidate drivers of transcriptomic state transitions via gallery-native reverse perturbation

**DOI:** 10.64898/2026.08.19.745674

**Authors:** Shiyang Liang, Cheng Yang, Jingjie Wang, Yunqi Li

**Affiliations:** Department of Gastroenterology, The No. 944 Hospital of Joint Logistic Support Force of PLA, Xiongguan Road, Jiuquan, 735000, China.; Department of Oncology, Air Force Medical Center, PLA, The Fourth Military Medical University, Beijing, 100142, China.; Department of Gastroenterology, Tangdu Hospital, The Fourth Military Medical University, Xi’an, 710038, China.; Department of Gastroenterology, The First Medical Center, Chinese PLA General Hospital, Beijing, 100853, China.

**Keywords:** RevPert, reverse perturbation, transcriptomic state transition, candidate driver, Perturb-seq, expression gallery, drug resistance

## Abstract

Cellular state transitions underlie adaptation, ageing and disease, yet prioritizing catalogued genetic perturbations whose expression signatures match an observed transcriptomic shift remains difficult. Most models predict phenotype from a nominated intervention, whereas genetic inverse benchmarks are largely restricted to within-screen identity recovery. Here we introduce **RevPert**, a gallery-native reverse perturbation model that predicts candidate drivers from a fixed genetic catalog for a query contrast **Δ*Y*** ^⋆^ **= *Y_B_ −Y_A_*** by combining signed Pearson connectivity with a learned residual. Across Replogle Essential Perturbseq (four lines) and LINCS-KO screens (ten lines), RevPert recovered held-out interventions at leading performance relative to matched baselines. Applied to public drug-resistance contrasts in HCC and CML, dual-arm prediction placed pre-specified disease anchors far higher on the expected arms than predicting from differential-expression magnitude alone (Essential residual model for HCC; a transductive GWPS residual for CML). RevPert therefore couples within-screen reverse prediction to a screen-external signed-geometry check; the latter calibrates literature anchors and is not claimed as held-out recovery.

## 1 Introduction

Cells move between consequential states during adaptation, ageing and disease, including drug resistance, senescence and malignant progression [1, 2]. These transitions can be represented by a transcriptomic contrast between an initial state *A* and a later or desired state *B*. Although differential-expression analysis identifies transcripts associated with the two states, it does not answer the intervention question: which catalogued genetic interventions are predicted to match, or reverse, the observed transcriptomic state transition? Such candidate drivers are useful because they turn an observed A-to-B expression shift into a shortlist of catalogued genetic hypotheses for follow-up, rather than a descriptive DEG list alone [3, 4]. Drug resistance is a natural external stress test of this idea: resistant and parental (or tissue) states define a clinically meaningful Δ*Y* ^⋆^, yet predicting candidates by differential-expression magnitude often fails to prioritize genes already implicated by orthogonal biology [5–7].

Pooled CRISPR screens with molecular readouts have created increasingly rich maps from genetic perturbations to expression responses [8–13]. Most computational work uses these resources for the forward task, predicting an expression response from a nominated perturbation [14]. Graph-based, foundation-model and knowledge-graph approaches have expanded this capability [15–17], while generative and transport-based models provide complementary formulations [18–21]. Recent cross-modal work, notably UniPert–G2CP, further bridges genetic screens and chemical phenotype prediction [22]. Together, these advances make perturbation profiles available as a reusable resource, yet none directly predicts which catalogued genetic interventions match an observed state transition.

Signature-to-intervention retrieval is well established for chemical perturbations through Connectivity Map, L1000 and LINCS [3, 4, 23–25]. For genetic perturbations, forward predictors can be repurposed as galleries for reverse matching, and CRISPR-GEM and PDGrapher place intervention recovery near the centre of their objectives [15, 16, 26, 27]. However, genetic inverse evaluation remains largely confined to held-out identities within the same screen. True screen-external use—predicting potential driver genes of an independently acquired state transition such as drug resistance—is rarely tested as a primary readout.

Here we define **reverse perturbation** as predicting candidate drivers from a fixed genetic catalog for a query contrast Δ*Y* ^⋆^ = *Y*_B_ − *Y*_A_ (Fig. 1). RevPert scores the query against predicted or observed knockout profiles by combining signed Pearson connectivity with a learned residual similarity, returning loss-of-function phenocopies and activation-like matches as a directional shortlist, not as experimentally established causal regulators. We audit this interface by held-out identity recovery across four Replogle Essential Perturb-seq and ten LINCS-KO cell-line datasets, retaining dataset-specific leakage-aware partitions [12, 15, 27]. We then apply the same dual-arm prediction interface to public HCC and CML resistance contrasts, assessing whether pre-specified anchors receive directionally appropriate ranks relative to differential-expression magnitude [5–7]. Together, these analyses present RevPert as a reverse model for predicting candidate drivers of transcriptomic state transitions: audited by within-screen identity recovery, and illustrated externally on drug resistance.

**Fig. 1.**
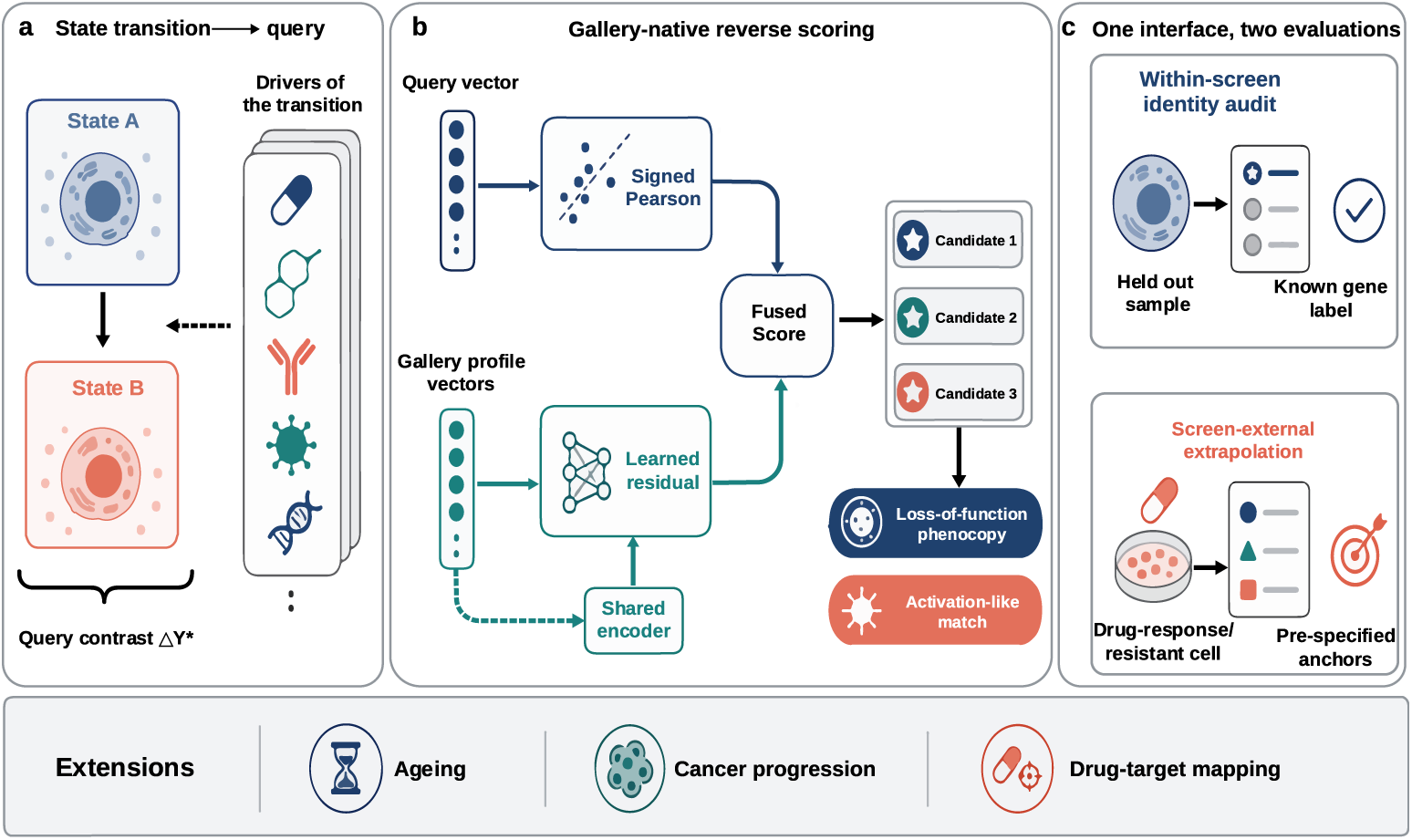
RevPert overview. **a**, Multiple upstream drivers of a cellular-state transition produce the query contrast Δ*Y* ^⋆^ = *Y_B_ −Y_A_*, which is compared with a genetic knockout expression gallery 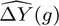 representing catalog genes. **b**, Gallery-native reverse scoring combines signed Pearson connectivity with a learned residual similarity derived from a shared profile encoder. Their fused score predicts candidate perturbations as loss-of-function phenocopies or activation-like matches. **c**, The same prediction interface is evaluated in two settings: within-screen held-out identity recovery, using known gene labels, and screen-external extrapolation to drug-resistance contrasts, using pre-specified biological anchors. The framework can be extended to ageing, cancer progression and drug–target mapping.

## 2 Results

### 2.1 A shared reverse-perturbation method across Essential and LINCS-KO

We formalized reverse perturbation as predicting catalog knockouts for a transcriptomic query against an expression gallery (Fig. 1). Identity recovery tests whether a held-out knockout can be retrieved from its observed control-subtracted response Δ*Y*. Primary metrics are median rank (lower is better) and Recall@10; PDGrapher-style partial% and nDCG are retained as Supplementary diagnostics.

Essential used Replogle Essential Perturb-seq in HepG2, K562, RPE1 and Jurkat [12, 15]. Queries were restricted to test knockouts present in both the observed response table and a shared predicted expression gallery 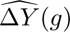 built by a companion linear progressive-stack forward model (hereafter the linear predicted gallery; Methods) [28]. RevPert was trained with fused InfoNCE on training knockouts only, learning a residual on top of signed Pearson gallery connectivity [29]. PCA compression was fit on training profiles only, and held-out observed responses were never used as gallery vectors. CMap baselines used the same mean-rank up/down enrichment against each resource’s matched expression gallery [3, 23, 28]. Matching observed responses to observed responses yielded median rank 1 on all four lines and was retained as an oracle upper bound in Supplementary Information only.

The same identity-recovery protocol was applied to ten LINCS-KO cell lines [27]. Gallery baselines, RevPert trained within each fold and the official PDGrapher model were scored under the same median-rank and Recall@10 definitions (Fig. 1; Fig. 2d; Supplementary Tables S9 and S10). Partition protocols for Essential and LINCS-KO are summarized in Supplementary Note S1. Beyond within-screen identity recovery, we reuse the same dual-arm prediction interface on public drug-resistance signatures as a screen-external stress test with literature anchors rather than single-gene labels (Essential residual model for HCC; a transductive GWPS residual for CML; detailed below).

**Fig. 2.**
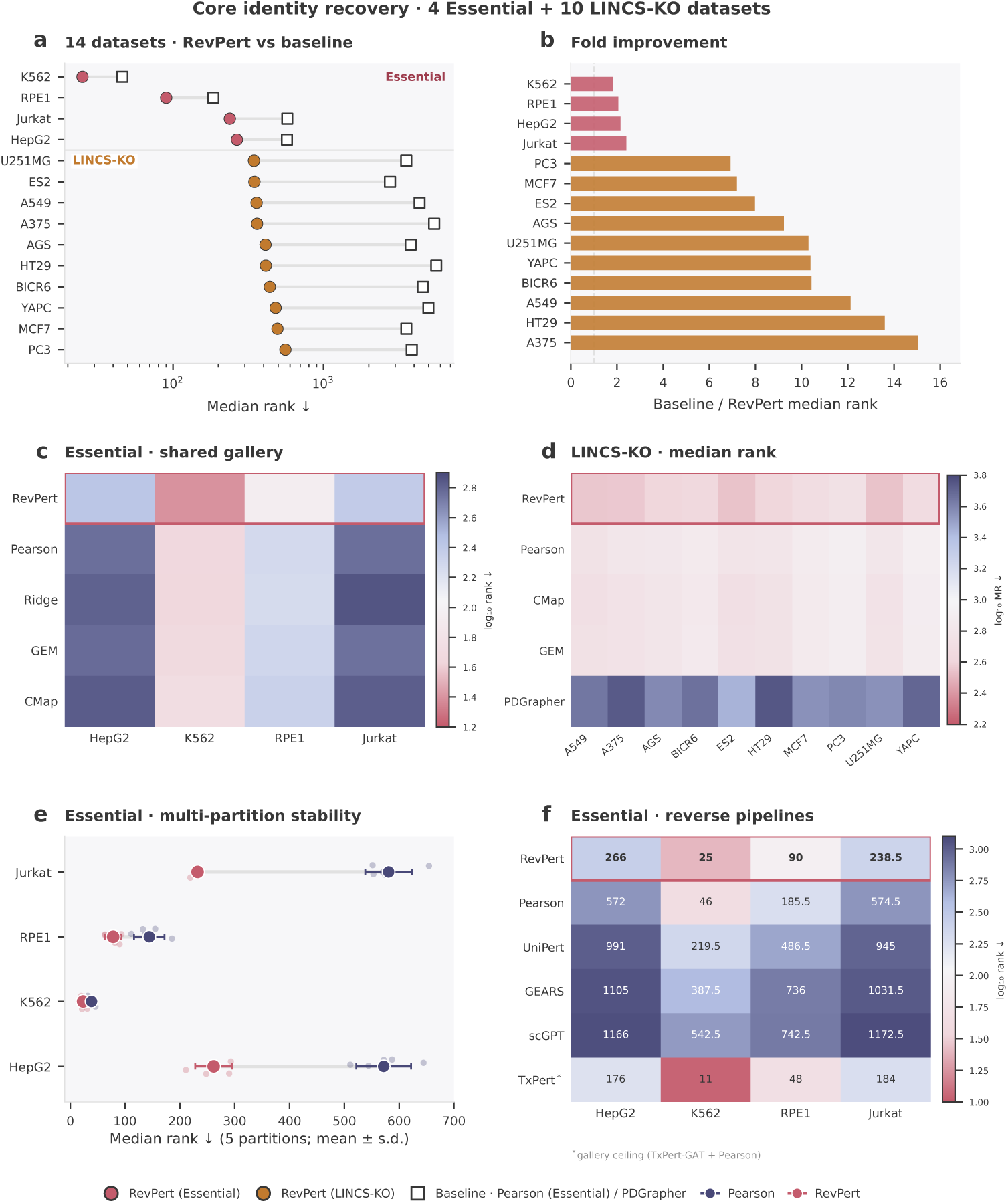
Main identity-recovery results across fourteen datasets. Four Essential and ten LINCS-KO lines under one reverse-perturbation task (not a pooled leaderboard). **a**, RevPert median rank versus the dataset-specific primary baseline (Pearson on Essential; official PDGrapher); RevPert best in 4/4 and 10/10 comparisons. **b**, Fold improvement (baseline / RevPert). **c**, Essential shared-gallery scorers (log_10_ median-rank heatmap). **d**, LINCS-KO mean median rank (ten lines *×* five folds). **e**, Essential multi-partition stability (five held-out partitions; mean*±*s.d.). **f**, End-to-end Essential reverse recipes (median rank): each method’s predicted gallery scored by Pearson, except RevPert (linear gallery + residual fusion); TxPert^∗^ is the within-line gallery ceiling. Numeric tables: Supplementary Tables S4, S12 and S13 (Supplementary Notes S2 and S4).

### 2.2 RevPert improves held-out identity recovery on Essential Perturb-seq

On the primary Essential held-out partition, RevPert achieved the best median rank among scorers on the shared linear predicted gallery in all four lines (Fig. 2a–c; Supplementary Table S4). Relative to Pearson matching on that gallery, median ranks improved by 1.8–2.4-fold (for example, HepG2 572 to 266), and Recall@10 rose by 1.5–3.4-fold (for example, HepG2 3.7% to 12.5%). Under hard within-screen labels, this Recall@10 gain is the operational candidate-shortlist readout: more true interventions fall inside a Top-10 follow-up list. Paired Wilcoxon signed-rank tests on per-query ranks rejected equality in favour of RevPert on every line (all *P <* 10^−18^), with bootstrap 95% confidence intervals on the median-rank difference excluding zero (Supplementary Table S6). The same ordering held across five independently drawn Essential partitions (20/20 line-partition comparisons; Fig. 2e; Supplementary Table S5). End-to-end reverse recipes built from GEARS, scGPT and UniPert galleries trail RevPert on every Essential line, whereas within-line TxPert-reverse is the absolute identity ceiling on that board (Fig. 2f; Supplementary Table S12) [15–17, 22].

GEM (DEG-subspace correlation) and CMap connectivity performed close to or worse than whole-profile Pearson matching on the predicted gallery (Fig. 2c; Supplementary Table S4). Ridge inversion from expression shifts to atlas coordinates was competitive on K562 but degraded on HepG2 and Jurkat. Matching top-magnitude genes to knockout sequence embeddings was a weak prior (median ranks 504–1220) and did not approach fair gallery methods. The observed-to-observed Pearson oracle recovered median rank 1 on all four lines (Supplementary Table S7). Additional diagnostics (Recall@1, Recall@100, mean reciprocal rank) reinforced the same RevPert *>* Pearson ordering, with the single exception of Jurkat Recall@1, where Pearson was marginally higher (1.35% versus 1.01%). Simple signature heuristics therefore did not close the RevPert gap under matched catalogs and queries.

### 2.3 RevPert improves held-out identity recovery on LINCS-KO screens

We next tested whether the reverse-perturbation method transfers beyond Essential Perturb-seq. On ten LINCS-KO cell lines (five folds), we scored Pearson, CMap and GEM gallery matching, RevPert trained within each fold and the official PDGrapher model under the same median-rank and Recall@10 definitions (Fig. 2d; Supplementary Tables S9 and S10). A protocol fairness audit confirmed matched splits, gene-axis restriction, Δ*Y* definition and prediction metrics, while noting that RevPert and PDGrapher remain different model classes (Supplementary Table S8). RevPert mean median ranks ranged from 345 to 560, lower than official PDGrapher in all ten lines (approximately 7–15×). Official PDGrapher mean median ranks remained in the thousands (approximately 2.8 × 10^3^–5.6 × 10^3^). Recall@10 followed the same ordering: RevPert improved over Pearson (4.14% versus 2.18%) and official PDGrapher (1.64%). We therefore treat median rank as the primary genetic recovery metric and Recall@10 as the candidate-shortlist diagnostic. Secondary diagnostics (Recall@1, partial%, nDCG) followed the same pattern (Supplementary Table S9). Per-line summaries are listed in Supplementary Table S10; these diagnostics were not used to redefine the identity-recovery main panel. Essential and LINCS-KO screens remain parallel evaluations of one method rather than one pooled prediction table. Catalog and query scales are reported in Supplementary Table S2. Partition-to-partition variation on Essential was modest relative to the RevPert–Pearson gap (Fig. 2e), confirming that the primary leaderboard is not a single-split artefact.

### 2.4 RevPert recovers signed resistance anchors across HCC and CML contrasts

Identity recovery validates catalog prediction under matched queries. We next asked whether the same fused RevPert scorer preserves signed connectivity on public resistance contrasts Δ*Y* ^⋆^: do disease-associated genes land near the top on the expected dual arm, and higher than under DEG-magnitude prediction [3, 4]? We focused on HCC under sorafenib and CML under imatinib (Table 1; Fig. 3). Signed Pearson dual-arm ranks are the geometry reference; fused RevPert is the scorer in Figure 3 and Table 1, with Pearson and learn-only ranks in Supplementary Table S16. This is a literature-anchored stress test of signed reverse prediction, not a hard-label Top-*k* contest against Pearson.

**Fig. 3.**
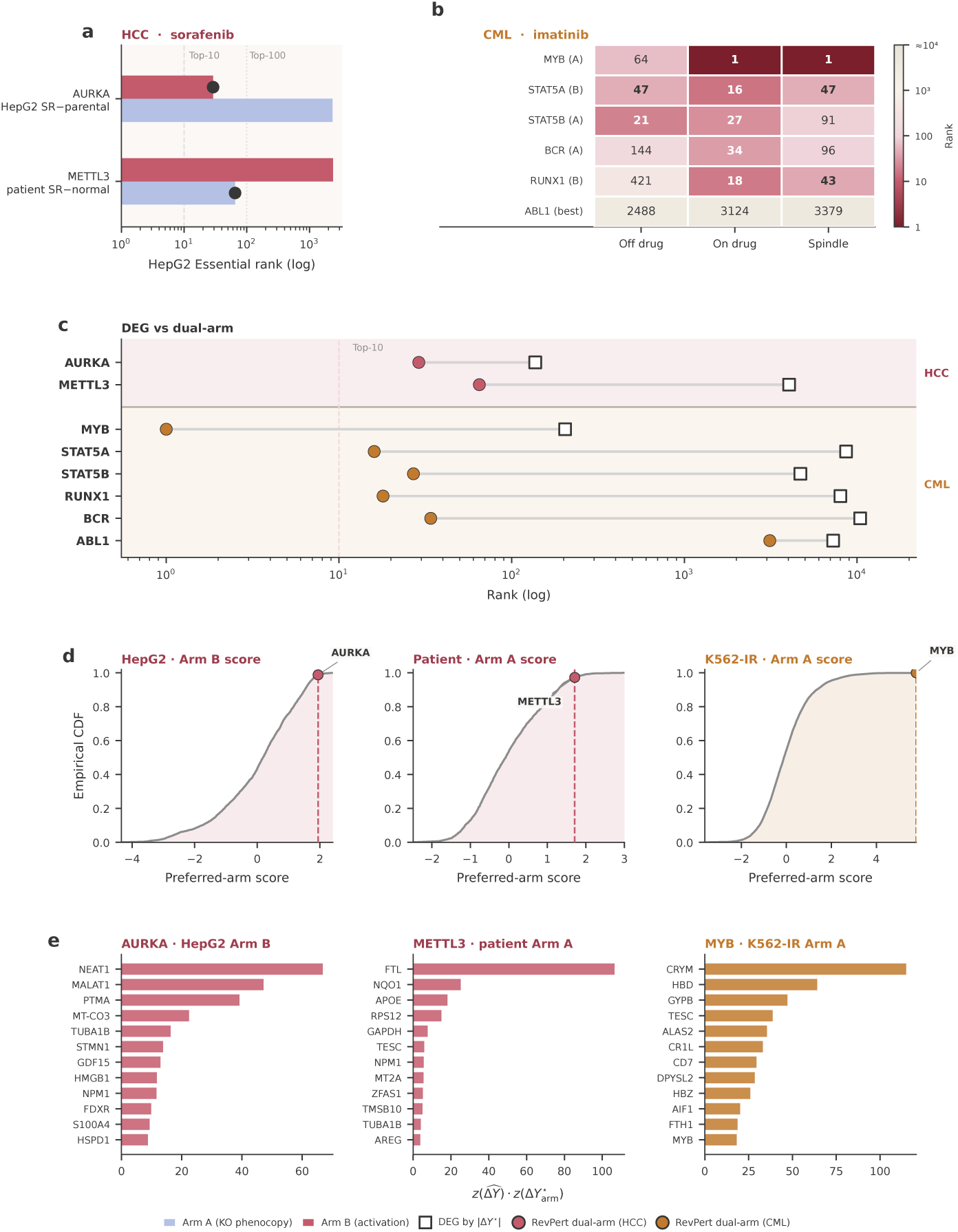
Resistance proving ground for two diseases. All panels use fused RevPert dual-arm scoring. **a**, HCC / sorafenib on HepG2 Essential; black dots mark preferred-arm ranks (AURKA 29; METTL3 65). **b**, CML / imatinib on K562 GWPS across three K562-IR contrasts (row tags A/B = preferred arm). **c**, Same anchors: squares, DEG rank by *|*Δ*Y* ^⋆^*|*; circles, RevPert preferred-arm rank (left better). **d**, Empirical CDFs of preferred-arm scores; literature calibrators sit in the upper tail. **e**, Top gene-level drivers of the gallery–query Pearson match preserved by RevPert. Pearson dual-arm ranks (AURKA 4; METTL3 6 on HCC) are in Table 1 and Supplementary Table S16.

**Table 1.** Two-disease resistance proving ground. Acquisition Δ*Y* ^⋆^ is resistant (or resistant tissue) minus the stated control. Headline ranks are fused **RevPert** preferred-arm ranks (Figure 3). Pearson dual-arm ranks in parentheses are reported in full as the *signed-geometry ceiling* on the same gallery (sharper on HCC; nearly coincident with RevPert on CML). DEG ranks are by *|*Δ*Y* ^⋆^*|* on the same signature.

| Disease | Drug / GEO | Contrast | Gallery | Headline ranks |
| --- | --- | --- | --- | --- |
| HCC | sorafenib<br>GSE322742 | / HepG2 SR – parental | Essential | AURKA Arm B **29** (Pearson 4; DEG $\sim 137$) |
| HCC | sorafenib<br>GSE143233 | / Patient SR – normal | Essential | METTL3 Arm A **65** (Pearson 6; DEG $\sim 4040$) |
| CML | imatinib<br>GSE120932 | / K562-IR – parental<br>($\times 3$) | GWPS | MYB **1**, STAT5A 16, RUNX1 18, STAT5B 27, BCR 34 (Pearson 1/13/20/25/36) |

Connectivity was scored by fused RevPert on each catalog (Figure 3); Pearson correlation *s* between each predicted knockout profile and the query remains the geometry reference. Arm A predicts knockouts by phenocopy of the query shift; Arm B predicts by the opposite signed match (activation heuristic). HCC used the HepG2 Essential gallery (*n* = 2372); CML used K562 GWPS (*n* = 9866) because classic CML drivers are largely absent from K562 Essential [12].

#### 2.4.1 Hepatocellular carcinoma (sorafenib)

Two sorafenib datasets probe opposite signed biology on the same HepG2 Essential gallery (Fig. 3a). Under fused RevPert, GSE322742 (HepG2 sorafenib-resistant versus parental) [5] places AURKA near last on Arm A (2331) but **29** on Arm B (DEG magnitude ∼137; Fig. 3a,c). On GSE143233 (patient sorafenib-resistant HCC versus normal liver) [6], METTL3 is **65** on Arm A and near last on Arm B (DEG ∼4040), consistent with loss-of-function evidence. Pearson geometry is sharper (AURKA Arm B 4; METTL3 Arm A 6); RevPert keeps both calibrators inside the Top-100 while trading some sharpness for the residual objective (Table 1). Preferred-arm score ECDFs and gene-level *z* × *z* drivers place both calibrators in the upper tail (Fig. 3d,e). Arm Top-50 versus magnitude-DEG Top-50 Jaccard was 0 under Pearson geometry, and a sign-flip null gave Top-50 overlap 0 (Supplementary Table S14).

#### 2.4.2 Chronic myeloid leukemia (imatinib)

GSE120932 compares parental K562 with imatinib-resistant K562-IR under three culture contrasts [7]. Associated CML pathway genes were predicted near the top of the GWPS catalog on the preferred arm and agreed across contrasts (Fig. 3b; Supplementary Table S15) [7, 30]. Under the on-drug contrast, MYB was Arm A **1**, STAT5A Arm B 16, STAT5B Arm A 27, RUNX1 Arm B 18 and BCR Arm A 34, with ABL1 mid-catalog (Pearson 1/13/25/20/36); DEG ranks were far worse (Fig. 3c). Sign-flip and gene-axis nulls under Pearson geometry keep the five pathway anchors extreme while ABL1 is not (Supplementary Fig. S1; Supplementary Table S14); fused RevPert ranks on the same contrast are in Supplementary Table S16. MYB sits in the extreme Arm A score tail, with match-driving genes dominated by erythroid/hemoglobin-linked features (Fig. 3d,e).

In both diseases, RevPert dual-arm prediction beat DEG-magnitude prediction (Fig. 3c), with Pearson the sharper geometry reference on the same catalogs. How residual fusion trades identity gains against that geometry is taken up next (Fig. 4). Catalog coverage for the main proving-ground signatures is summarized in Supplementary Table S3.

**Fig. 4.**
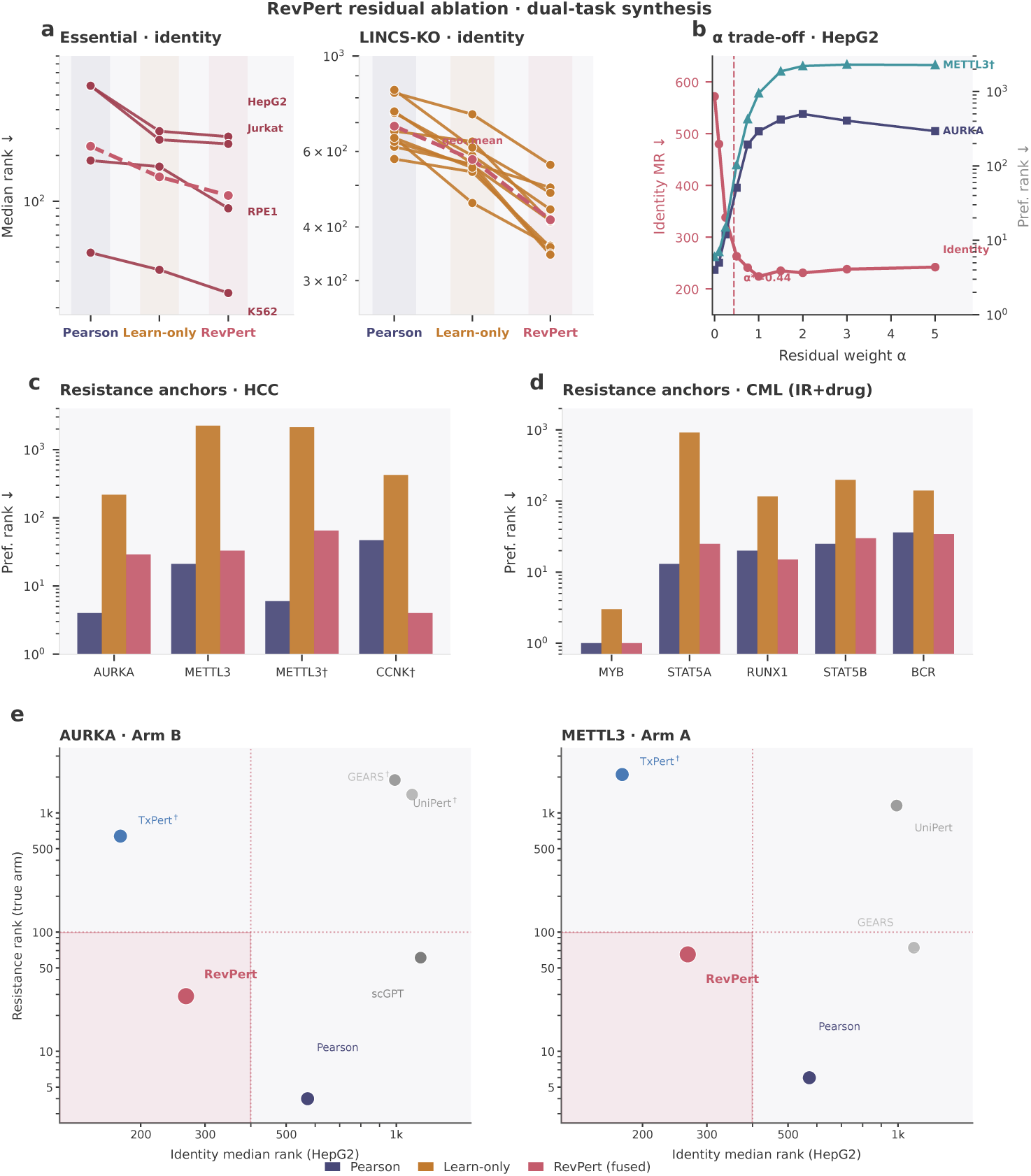
RevPert residual ablation and identity–resistance synthesis. **a**, Identity median-rank trajectories for Pearson, learn-only and RevPert (Essential left, seed 1; LINCS-KO right, fold means; log scale). Dashed pink lines, within-panel geometric means. **b**, HepG2 *α* scan: identity median rank (circles) versus preferred-arm ranks of AURKA (squares) and METTL3 (triangles); dashed line, training-selected *α*^⋆^. **c**, Preferred-arm ranks of HCC resistance anchors under the three scorers (*†*, patient SR*−*normal). **d**, Preferred-arm ranks of CML IR+drug anchors on K562 GWPS. **e**, Identity versus true-arm resistance rank for AURKA (Arm B) and METTL3 (Arm A); shaded region, identity *<* 400 and resistance rank *<* 100. *†*, preferred arm disagrees with literature expectation. Pipeline identity ranks are Figure 2f.

### 2.5 Residual fusion retains identity gains without collapsing resistance anchors

Having established identity recovery and the resistance proving ground, we asked which components of RevPert are necessary (Fig. 4). Across all 14 identity datasets, Pearson matching remained weak for identity, whereas learn-only improved median ranks but still trailed fused RevPert (Fig. 4a; best in 4/4 Essential and 10/10 LINCS-KO). An *α* scan on HepG2 shows the trade-off: larger residual weight lowers identity median rank until preferred-arm ranks of AURKA and METTL3 rise sharply outside the small-residual regime (Fig. 4b; *α*^⋆^ = 0.44). Learn-only displaces HCC and several CML anchors by orders of magnitude, whereas RevPert holds HCC calibrators inside the Top-100 and tracks Pearson on GWPS CML anchors (Fig. 4c,d). Residual fusion therefore couples identity discrimination to signed connectivity. On the identity–resistance board, only RevPert clears both operational gates (Fig. 4e): Pearson is the sharper signed-geometry reference, and within-line TxPert-GAT is the held-out identity ceiling (Fig. 2f) yet fails both HCC arm polarities; GEARS, UniPert and scGPT miss at least one gate. Gallery diagnostics link part of the TxPert resistance failure to collapsed within-line spectra (Supplementary Table S13). Absolute identity and signed resistance are therefore separate evaluations; RevPert is the joint claim on a deployable linear gallery, not a bid to beat TxPert absolute identity.

## 3 Discussion

Under matched Essential and LINCS-KO splits, residual RevPert outperformed Pearson gallery matching and led official PDGrapher on median rank [27]. Ablations show that learn-only matching improves identity but collapses resistance anchors, whereas residual fusion retains both (Fig. 4). Within-line TxPert-reverse is the held-out identity ceiling on Essential (Fig. 2f) but does not redefine the residual reverse claim (Supplementary Table S12) [15–17, 22]. UniPert–G2CP advances forward, cross-modal perturbation-to-phenotype transfer; RevPert addresses the complementary phenotype-to-catalog prediction problem. These conclusions are falsifiable on frozen Essential and LINCS-KO prediction tables.

A deployable reverse scorer should preserve signed connectivity while learning discriminative residuals, rather than discarding Pearson geometry for unconstrained retrieval [28]. Gallery fidelity still matters for CMap-style methods, so forward and reverse evaluation remain coupled but not interchangeable [3, 4]. We therefore report two ceilings: within-line TxPert-reverse for held-out identity recovery (Fig. 2f) and Pearson dual-arm matching for signed geometry on the linear gallery (Table 1) [5–7]. RevPert stays near the front on both under one gallery, whereas the TxPert identity ceiling fails HCC arm calibrations (Fig. 4e; Supplementary Table S13). Unsigned Top lists on a knockout gallery remain mechanistically ambiguous without signed dual-arm readout.

Limitations are largely data-bound. Identity recovery and therapeutic/resistance prediction are separate tasks; only the former is claimed as a held-out prediction evaluation. The resistance proving ground is an out-of-distribution use of Perturb-seq knockout galleries for signed calibration under public acquisition signatures [12]. A gene can be predicted only if it is in the scored catalog; when Essential coverage is sparse we used GWPS [12, 13]. Cross-context encoder transfer degrades for all Supplementary Table S4 scorers, especially residual and learn-only encoders, motivating lineage-matched catalogs (Supplementary Tables S11, S12; Supplementary Note S4). Identity-recovery median ranks near 10^2^ and proving-ground Top ranks therefore answer different questions. The CML arm is transductive (residual fitted over the full GWPS catalog), so it is a signed-geometry check rather than generalisation evidence. Tissue bulk mixes composition with cell-autonomous programmes; CRISPR-GEM phenotype simulators remain outside the fair Essential comparison [26].

As Perturb-seq atlases expand, the same reverse interface can be retrained on broader catalogs [12, 13]. Downstream of reverse prediction (outside the present evaluation), shortlists can feed drug–target models such as TAPB or DrugCLIP and focused wet-lab assays [31, 32]. Beyond resistance, the same prediction task applies to other state transitions when matched catalogs exist (for example colorectal progression or senescence programmes) [1, 2]; free cross-context encoder transfer is not assumed (Supplementary Note S4).

## 4 Methods

### 4.1 Overview and estimand

RevPert formulates reverse perturbation as predicting candidate drivers from a fixed intervention catalog given a transcriptomic query. Identity recovery tests whether a held-out knockout can be recovered from its observed control-subtracted response under matched splits and catalogs. Public resistance signatures serve as a literature-anchored proving ground for the same gallery scoring: pre-specified signed anchors should be predicted on the expected dual arm. Two datasets are evaluated for identity recovery without pooling into a single leaderboard: Replogle Essential Perturb-seq in four cell lines, and LINCS L1000 genetic knockout screens (LINCS-KO) in ten cell lines [12, 27]. Primary metrics are median rank (lower is better) and Recall@10. Additional diagnostics are defined below and reported in Supplementary Information unless otherwise noted.

### 4.2 Datasets and processing

#### Essential — Replogle Essential Perturb-seq

##### Source

Genome-scale Essential Perturb-seq profiles for HepG2, K562, RPE1 and Jurkat were obtained from Replogle et al. [12] and processed in the GEARS-compatible single-cell format used by the companion forward-prediction stack [15]. After single-gene filtering and gallery intersection, catalog sizes were 2372 (HepG2), 1086 (K562), 1534 (RPE1) and 2372 (Jurkat) knockouts, with gene axes of 4,261–5,000 features aligned to the corresponding forward gallery.

##### Pseudo-bulk ΔY

Cells were grouped by perturbation condition within each cell line. The control baseline was the mean expression of all unperturbed control cells. For each single-gene knockout condition, pseudo-bulk Δ*Y* was the condition mean minus the control baseline. Multi-gene combinatorial conditions and the control condition itself were excluded, yielding one Δ*Y* vector per single-gene knockout on the shared gene axis.

##### Held-out queries

Identity-recovery queries used the primary held-out knockout partition for each Essential line (Supplementary Note S1). Only knockouts present in both the observed Δ*Y* table and the predicted expression gallery were retained for prediction.

##### Forward gallery for Essential

The default Essential catalog is a predicted expression gallery from a companion linear progressive-stack model trained under leave-one-line-out stacking (linear predicted gallery) [28]. Per line, TruncatedSVD on all pseudo-bulk Δ*Y* profiles yielded a 10-dimensional knockout embedding; the other three lines’ 10-d embeddings were concatenated to form a 30-dimensional atlas coordinate *P* for the held-out line. The forward model predicted absolute expression from *P*; predicted Δ*Y* for gallery matching was the absolute prediction minus the matched control profile on the same gene axis. Queries for Pearson, CMap and GEM were aligned to this gallery gene list; genes absent from the gallery were omitted from the correlation.

#### LINCS-KO — LINCS L1000 genetic knockout screens

##### Source

These CRISPR knockout screens were compiled by Gonzalez et al. [27] from LINCS Level 3 plate-normalized expression [23] intersected with BioGRID protein–protein interactions [33]. Unlike Essential, LINCS-KO uses plate-based LINCS L1000 expression profiles rather than single-cell Perturb-seq. Ten cell lines met the published coverage filter (≥4,000 unique genetic perturbations): A549, A375, AGS, BICR6, ES2, HT29, MCF7, PC3, U251MG and YAPC. After gene–PPI intersection, the graph comprised 10,716 nodes with approximately 1.5 × 10^5^ undirected edges; samples whose target gene fell outside the PPI were removed. Each sample provides a baseline expression state, a post-knockout state and an intervention indicator on the graph.

##### Reverse ΔY and evaluation folds

Per-sample Δ*Y* was defined as treated minus diseased (PDGrapher terms for post-knockout minus baseline) on the 10,716-gene axis. Train-fold galleries aggregated mean Δ*Y* by intervention gene within the training indices of each fold. Within each official LINCS-KO fold, queries and catalogs were restricted to genes available to all compared methods before computing median rank and Recall@10 (Supplementary Note S1). Official PDGrapher models were scored under the same identity-recovery prediction interface without modifying the published training recipe beyond runtime settings needed for stable evaluation. Matched and unmatched protocol dimensions are summarized in Supplementary Table S8.

#### External resistance and disease signatures

Primary HCC proving-ground queries were scored against the HepG2 Essential linear predicted gallery after intersecting Δ*Y* ^⋆^ with the gallery gene axis (Pearson correlation requires ≥ 10 finite overlapping genes). Figure 3 and Table 1 report fused RevPert dual-arm ranks as the primary proving-ground readout. Pearson and learn-only ranks for the same queries are in Supplementary Table S16. Primary calibrators were:

- GSE322742 HepG2 sorafenib-resistant versus parental RNA-seq [5]: Δ*Y* ^⋆^ = mean log_2_(FPKM+ 1)_SR_ −mean log_2_(FPKM+ 1)_parental_ (two replicates per group); Ensembl identifiers were mapped to gene symbols and duplicate symbols averaged.
- GSE143233 patient sorafenib-resistant HCC versus normal liver [6]: Δ*Y* ^⋆^ = log_2_(FPKM+1)_SR_ −log_2_(FPKM+1)_normal_ using the GEO-released cohort-averaged columns.

Blood-lineage dual-arm queries used the Replogle K562 GWPS observed pseudobulk knockout gallery (*n* = 9866 interventions after control filtering) [12] as the primary catalog. The K562 Essential linear predicted gallery was retained only as a coverage control (classic CML/AML drivers were largely absent). GSE120932 parental K562 versus imatinib-resistant K562-IR microarray contrasts [7] defined three acquisition signatures as resistant-group mean minus parental mean (IR cultured off drug; IR maintained in imatinib; spindle-shaped adherent IR). All three CML contrasts were scored with Pearson, learn-only and fused RevPert after training a GWPS gallery-native residual model on noise-augmented observed queries (Supplementary Table S16). This GWPS model is deliberately transductive: the residual encoder uses the full observed knockout catalog (*n* = 9866) as gallery vectors, and noise-augmented validation queries are perturbed copies of catalog entries, so validation identity recovery saturates (median rank 1, MRR 1.00) and cannot discriminate between checkpoints. Epoch selection therefore falls back to the fused validation objective, and the deployed residual weight was set by leave-self-out retune (*α* = 0.1), small enough that fused CML ranks track Pearson closely (score correlation *>*0.99; identical preferred-arm Top-10 sets). The GWPS arm is consequently reported as a signed-geometry stress test of a small learned residual, not as evidence that the learned component generalizes on this catalog; all quantitative identity claims rest on Essential and LINCS-KO.

#### Gene sequence control

Human UniProt protein sequences for Essential-panel gene symbols [34] were embedded with ESM2 (650M parameter model; mean-pooled residue representations, 1280-d) [35]. These embeddings were used only for the sequence-only ESM-mean control in Supplementary Table S7 and are not inputs to RevPert.

#### Deposition

Split lists, frozen identity-recovery prediction tables, LINCS-KO fairness audit tables and Essential paired-rank exports are available with the analysis repository as described under Data availability and Code availability.

### 4.3 RevPert gallery-native reverse model

RevPert is a residual prediction model that scores a query expression shift against a fixed expression knockout gallery. It is intentionally not a gene-sequence or graph encoder: catalog objects are expression profiles 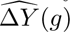 (predicted for Essential; train-fold means for LINCS-KO; observed for the GWPS proving-ground catalog), and the learnable component is a profile encoder that adds a residual to signed Pearson connectivity.

#### Inputs and PCA

Queries and gallery profiles are control-subtracted expression vectors on a shared gene axis. A PCA compressor (*d*_PCA_ = 256) is fit on training profiles only (observed training queries plus the corresponding training gallery profiles) and applied to both queries and gallery vectors at train and test time. Held-out observed responses are never used to fit PCA.

#### Shared profile encoder

Let *P* (·) denote the PCA map. A shared multilayer perceptron *f*_θ_ embeds *P* (*q*) and 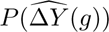 into a common 128-dimensional *ℓ*_2_-normalized space (two hidden layers of width 512, GELU activations, dropout 0.1). Learned similarity is

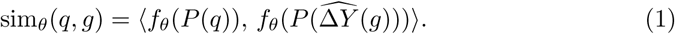

Query and gallery towers share weights, so the model learns a profile geometry rather than separate query/gallery spaces.

#### Residual fusion

The deployed score is a per-query *z*-scored residual fusion of Pearson connectivity and learned similarity:

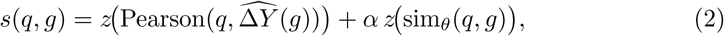

with *α* = softplus(*β*) and *β* initialized at −1 (*α* ≈ 0.31) so that training begins near the Pearson geometry. Setting *α* = 0 recovers Pearson; *α* → ∞ recovers learn-only matching. Both arms of the resistance proving ground reuse the same score with *s* versus −*s*.

#### Training objective

Training maximizes a full-catalog InfoNCE loss on the fused logits so that the true gallery entry of each training query is recovered [29], with a mild Pearson-teacher regularizer on the learned tower (KL divergence toward a 50/50 mix of the hard identity label and a temperature-sharpened Pearson distribution over the catalog; weight 0.25). Optimizer defaults are AdamW (10^−3^, weight decay 10^−4^) with cosine learning-rate decay. Essential models use batch size 64 for 40 epochs; LINCS-KO fold models use batch size 256 for 25 epochs. Checkpoints and *α* are selected on validation fused identity recovery (validation MRR, with learned-score Spearman-to-Pearson as tie-break). Architecture and training defaults are summarized in Supplementary Note S6 (Supplementary Table S17).

#### Protocol by dataset

For both identity-recovery datasets, catalog entries are predicted or train-fold gallery profiles; held-out observed responses are never used as gallery vectors. RevPert is retrained independently on each Essential partition and each LINCS-KO fold (Supplementary Note S1). The screen-external K562 GWPS model differs: it is a transductive residual scorer over the full observed knockout catalog (*n* = 9866), trained on noise-augmented observed queries, with no held-out identity validation (Methods, External resistance and disease signatures).

### 4.4 Baselines and related methods

#### Fair Essential baselines

Pearson matching predicted catalog interventions by Pearson correlation between the query and the predicted gallery profile 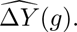 CMap scored each gallery profile by mean-rank enrichment of the query up- and down-regulated gene sets (top 100 genes by signed magnitude), using the same genetic gallery as Pearson and GEM rather than the Broad chemical Touchstone database [3, 23]. The identical CMap definition was used on Essential and LINCS-KO. GEM restricted correlation to a DEG subspace (top 200 genes by |Δ*Y* | in the query) as a lightweight control inspired by CRISPR-GEM, not a full phenotype simulator [26]. Ridge inversion fit a linear map from training Δ*Y* to atlas *P* (ridge penalty *α* = 1) and predicted catalog interventions by cosine similarity of the mapped query. ESM2 sequence-prototype matching (top 50 genes by magnitude) is reported only as a weak sequence prior (Supplementary Table S7). Observed-to-observed Pearson matching is Supplementary-only as an oracle upper bound (Supplementary Table S7).

#### End-to-end reverse pipelines

GEARS, scGPT, TxPert and UniPert were evaluated as complete reverse pipelines: each builds 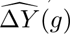 under the matched Essential seed-1 split, then predicts held-out identities by Pearson matching to that gallery (Fig. 2f; Supplementary Tables S12 and S13; Supplementary Note S4) [15–17, 22]. RevPert is residual fusion on the linear predicted gallery, not a scorer substituted onto those foreign catalogs as the primary claim. For UniPert-reverse, a published UniPert gene embedding was held fixed and a Ridge head mapped embeddings to Δ*Y* on training knockouts before Pearson matching. scGPT knockouts absent from the shared catalog received the worst rank. TxPert-reverse (within-line TxPert-GAT gallery + Pearson) is reported as a gallery-ceiling reference on the Essential identity board; residual RevPert retrained on foreign galleries and TxPert cross-cell leave-one-line-out remain Supplementary sensitivity checks (Supplementary Table S12). As a separate encoder-transfer diagnostic (Supplementary Note S4; Supplementary Tables S11 and S12), seed-1 Essential shared-gallery scorers (Supplementary Table S4) were applied across lines under a shared gene-axis alignment. Ridge uses the source *Y* → *P* map and source *P* atlas; expression-gallery scorers use the target linear predicted gallery reindexed to the source gene axis. This diagnostic motivates lineage-matched catalogs for screen-external analyses and is not part of the primary leaderboard. Knockouts absent from an incomplete gallery (scGPT) were assigned the worst possible rank so that coverage enters the metric.

#### LINCS-KO baselines

On LINCS-KO, Pearson, CMap and GEM gallery matching, RevPert and the official PDGrapher model were scored under the same median-rank and Recall@10 definitions; Essential uses the same CMap and GEM definitions on the shared linear predicted gallery. Full CRISPR-GEM phenotype-shift models remain outside the shared-gallery Essential comparison because their objectives differ from identity recovery under a shared predicted gallery [26].

### 4.5 Data analysis

#### Ranking interface

Each method returned a full ordering of catalog interventions for every query. Scoring rules follow Sections 4.3–4.4; all outputs were converted to full-catalog predictions before metric computation. Ties were broken by catalog index.

#### Primary and secondary metrics

Let *r*_i_ be the rank of the true intervention for query *i*, counting from 1. Primary metrics were the median of {*r*_i_} over test queries and Recall@10 (fraction of queries with *r*_i_ ≤ 10; Fig. 2). Screen-external proving-ground claims use preferred-arm ranks of pre-specified anchors rather than Recall@10, reported as fused RevPert dual-arm ranks in Figure 3 and Table 1, with Pearson geometry ranks alongside (Supplementary Table S16). As a gallery-swap diagnostic, the two HCC anchors were predicted by Pearson dual-arm connectivity on GEARS, scGPT, UniPert-ridge and TxPert Essential galleries alongside the linear predicted gallery (Fig. 4e; Supplementary Note S4; Supplementary Tables S12 and S13). The RevPert point on Figure 4e uses fused dual-arm ranks on the linear gallery (AURKA Arm B 29; METTL3 Arm A 65), not Pearson. This board separates absolute identity recovery from signed resistance calibration; it does not redefine the residual RevPert claim. For LINCS-KO five-fold experiments, we report fold-wise mean±s.d. of median rank and Recall@10, and the mean of cell-line means when summarizing ten lines. Catalog and query counts are in Supplementary Table S2; ranks were never pooled across Essential and LINCS-KO. Mean rank, Recall@1/@100/@1000, MRR, partial% and nDCG were computed as diagnostics. On single-true-target identity-recovery queries, partial% coincides with Recall@1. nDCG used the PDGrapher-style gain (1 − *r/N*)*/* log_2_(*r* + 1) normalised by the ideal single-hit discounted cumulative gain, where *N* is catalog size [27, 36]. These diagnostics were not used to redefine the main leaderboard (Supplementary Information).

#### Statistical analysis

The primary Essential contrast compared RevPert and Pearson predicted-gallery ranks on the same held-out query sets. On the primary partition, paired per-query ranks were exported, a one-sided Wilcoxon signed-rank test was applied (alternative: Pearson rank *>* RevPert rank), and a bootstrap 95% confidence interval (*B*=2000) was computed on the difference of medians (Supplementary Table S6). Multi-partition stability summarized mean±s.d. of median rank and Recall@10 over five independently drawn held-out partitions, each with a retrained RevPert model (Supplementary Note S1; Fig. 2e). LINCS-KO comparisons are reported as fold-averaged point estimates with fold s.d.; formal paired tests were not pooled across cell lines. LINCS-KO fairness relative to official PDGrapher is documented as a protocol checklist rather than a pooled *P* -value (Supplementary Table S8).

### 4.6 Signed resistance proving-ground scoring

#### Scoring

The primary HCC proving ground uses the HepG2 Essential gallery and dual-arm connectivity defined above. Connectivity was computed as Pearson correlation between each predicted knockout profile and the query signature on overlapping finite genes. Arm A predicts by phenocopy (*s*); Arm B predicts by the opposite signed match (−*s*), an activation heuristic. Literature anchors in the gallery (AURKA, METTL3) were reported on both arms as pre-specified checks. For the HCC residual ablation, both arms were rescored with Pearson, learn-only similarity and fused RevPert using the training-selected *α*. DEG controls used the Top-50 genes by absolute query shift; sign-flip nulls negated the query before rescoring and compared Top-50 lists by overlap and Jaccard index. For the three GSE120932 contrasts, empirical preferred-arm *P* values for pre-specified CML anchors were computed under Pearson dual-arm scoring with 500 gene-axis permutations of Δ*Y* ^⋆^ on the GWPS catalog (Supplementary Fig. S1; Supplementary Table S14). These nulls test signed Pearson geometry, not the fused GWPS residual. Blood-lineage extension reused Arm A/Arm B scoring on the K562 GWPS observed gallery for GSE120932, with Essential K562 retained only to document driver-coverage gaps. Neither setting yields validated drivers or therapeutic targets.

### 4.7 Software and reproducibility

Analyses used Python 3.10.14 with NumPy 2.2.6, SciPy 1.15.3 and pandas 2.3.3. RevPert training and inference used PyTorch 2.12 (CUDA 13.0) on a single NVIDIA RTX 5090 GPU. Essential partitions were drawn with integer seeds 1–5 under the GEARS simulation protocol (Supplementary Note S1). Network initialization and batch ordering are not separately seeded, so retraining reproduces the reported orderings and fold-level summaries rather than bitwise-identical rank tables. PDGrapher training and scoring followed the published repository recipe; only dataloader workers and batch size were adjusted for stable evaluation. Analysis code, data-processing scripts and frozen tables are available as described under Code availability and Data availability.

## 5 Data availability

Essential profiles are publicly released Replogle Essential Perturb-seq data [12, 15]. LINCS-KO profiles are from the screens released with Gonzalez et al. [27], derived from LINCS Level 3 [23]. Blood-lineage dual-arm analyses additionally use the Replogle K562 GWPS observed pseudobulk knockout gallery [12]. Held-out partition definitions are given in Supplementary Note S1. Proving-ground signatures are derived from the stated GEO accessions (GSE322742, GSE143233, GSE120932) [5–7]. Frozen identity-recovery prediction tables, Essential per-query ranks, LINCS-KO fairness audit tables and split lists are available with the analysis repository (https://github.com/gityuling/RevPert; see Code availability). Primary source datasets remain available from the cited public resources.

## 6 Code availability

Data-processing and analysis code, together with preflight reproduction scripts and frozen SI tables, are available at https://github.com/gityuling/RevPert under the MIT license.

## 7 Acknowledgements

This work was supported by the National Key R&D Program of China (2024YFF1206700).

## 8 Author contributions

S.L. conceived the study, developed RevPert, performed the analyses and wrote the manuscript. C.Y. proposed the resistance proving-ground analyses and revised the manuscript. J.W. revised the manuscript. Y.L. supervised the project and provided critical revision. All authors approved the final manuscript.

