## Supplementary Information for "RevPert: predicting candidate drivers of transcriptomic state transitions via gallery-native reverse perturbation"

Primary claims in the main text are supported by Supplementary Notes S2, S3 and S5. Supplementary Note S4 motivates lineage-matched catalogs; Supplementary Notes S1 and S6 record protocols and training defaults.

This supplement is organised as follows: Supplementary Note S1, protocols and dataset inventories; Supplementary Note S2, Essential identity-recovery expansions; Supplementary Note S3, LINCS-KO expansions; Supplementary Note S4, cross-context transfer and gallery diagnostics; Supplementary Note S5, resistance proving-ground supplements; Supplementary Note S6, model hyperparameters.

### S1 Protocols and dataset inventories

Identity recovery uses *two different partition protocols* on Essential and LINCS-KO; they are not interchangeable and are never pooled into one leaderboard.

##### *Essential.*

Train / validation / test lists are knockout-level partitions under the GEARS *simulation* protocol [1]: genes are held out as entire interventions rather than splitting cells within a knockout. A single protocol is used throughout; integers 1–5 are random seeds that draw five independent partitions under that same protocol (they are not GEARS model-training seeds and are not a second splitting method). The primary leaderboard uses seed 1; multi-partition stability retrains and rescores on seeds 1–5. Approximate seed-1 test sizes before catalog filters: HepG2 594, K562 272, RPE1 384, Jurkat 594; after restricting to knockouts present in both the observed table and the gallery, the scored query counts are 593, 272, 384 and 594 (Supplementary Table S2). Only knockouts present in both the observed $\Delta Y$ table and predicted expression gallery enter prediction.

##### *LINCS-KO.*

Partitions are the official within-line random five-fold splits released with Gonzalez et al. [2] (backward indices for reverse prediction). This is a sample-level fold protocol on LINCS-KO screens, distinct from Essential knockout-level simulation partitions. We do not use leave-cell-out chemical splits. Within each fold, queries and catalogs are restricted to genes available to all compared methods.

**Table S1 Partition protocols at a glance.**

| Dataset | Protocol | What varies |
|---|---|---|
| Essential | GEARS simulation (KO hold-out) | Seeds 1–5 = five draws of the same protocol |
| LINCS-KO | Official random 5-fold | Folds 1–5 within each cell line |

***Inventories.***

Catalog and query counts for identity recovery, plus sample sizes for the main proving-ground signatures. Essential profiles are Replogle Essential Perturb-seq [3]; LINCS-KO profiles are from the screens released with Gonzalez et al. [2]. Essential and LINCS-KO are never pooled into one prediction table.

**Table S2 Fourteen-line dataset inventory.** Catalog size and test-query counts used for identity recovery. Essential: primary held-out partition; LINCS-KO: fold-wise query counts (means reported in main tables use official five-fold summaries).

| Dataset | Cell line | Catalog $n$ | Test queries $n$ |
|---|---|---|---|
| Essential | HepG2 | 2372 | 593 |
| Essential | K562 | 1086 | 272 |
| Essential | RPE1 | 1534 | 384 |
| Essential | Jurkat | 2372 | 594 |
| LINCS-KO | A549 | 10716 | 4851 |
| LINCS-KO | A375 | 10716 | 4359 |
| LINCS-KO | AGS | 10716 | 4257 |
| LINCS-KO | BICR6 | 10716 | 4237 |
| LINCS-KO | ES2 | 10716 | 4742 |
| LINCS-KO | HT29 | 10716 | 4105 |
| LINCS-KO | MCF7 | 10716 | 3755 |
| LINCS-KO | PC3 | 10716 | 4246 |
| LINCS-KO | U251MG | 10716 | 5265 |
| LINCS-KO | YAPC | 10716 | 4027 |

**Table S3 External-signature inventory (main proving ground).** Sample sizes, gallery and role for the HCC and CML signatures used in the main text.

| Signature | Contrast | Samples | Gallery | Role |
|---|---|---|---|---|
| GSE322742 | HepG2 SR vs parental | 2 vs 2 | HepG2 Essential ($n$=2372) | Main proving ground |
| GSE143233 | Patient SR vs normal | 3 vs 3 (GEO avg.) | HepG2 Essential | Main proving ground |
| GSE120932 | K562-IR vs parental (3 contrasts) | microarray groups | K562 GWPS ($n$=9866) | Main proving ground |

### S2 Essential identity-recovery expansions

The Essential shared-gallery numeric leaderboard is Table S4 below; end-to-end reverse recipes are visualised in main-text Figure 2f, with gallery diagnostics in Supplementary Tables S12 and S13 (Supplementary Note S4; Figure 4e).

The tables below expand multi-partition stability, paired RevPert–Pearson tests and related diagnostics.

**Table S4 Essential identity-recovery leaderboard (primary partition).** Lower median rank is better. All scorers use one shared linear predicted gallery (primary fair comparison for RevPert). Boldface marks the best entry. Complements main-text Figure 2a–c; end-to-end reverse recipes versus GEARS-/scGPT-/TxPert-/UniPert-reverse are shown in main-text Figure 2f and Supplementary Tables S12 and S13 (Supplementary Note S4), and do not redefine this same-gallery leaderboard.

| Method | Metric | HepG2 | K562 | RPE1 | Jurkat |
|---|---|---|---|---|---|
| RevPert | Median rank | **266** | **25** | **90** | **238.5** |
| | Recall@10 (%) | **12.5** | **27.6** | **16.7** | **11.4** |
| Pearson (linear gallery) | Median rank | 572 | 46 | 185.5 | 574.5 |
| | Recall@10 (%) | 3.7 | 18.8 | 10.2 | 5.6 |
| Ridge $\Delta Y \to P$ | Median rank | 638 | 45.5 | 180 | 708 |
| | Recall@10 (%) | 5.6 | 21.3 | 12.5 | 5.1 |
| GEM | Median rank | 587 | 48.5 | 195 | 565.5 |
| | Recall@10 (%) | 4.0 | 18.8 | 11.2 | 5.7 |
| CMap | Median rank | 668 | 53.5 | 215.5 | 653 |
| | Recall@10 (%) | 3.7 | 21.0 | 9.9 | 4.9 |

**Table S5 Identity-recovery multi-partition stability (Essential).** Mean±s.d. of median rank and Recall@10 (%) over five held-out partitions under the GEARS simulation protocol (Supplementary Note S1). RevPert is retrained independently per partition; Pearson uses the same linear predicted gallery protocol.

| Method | Metric | HepG2 | K562 | RPE1 | Jurkat |
|---|---|---|---|---|---|
| RevPert | Median rank | $261.6 \pm 33.8$ | $23.4 \pm 4.8$ | $78.3 \pm 14.4$ | $232.2 \pm 8.2$ |
| | Recall@10 (%) | $12.9 \pm 0.7$ | $31.8 \pm 3.7$ | $19.6 \pm 2.8$ | $11.3 \pm 0.8$ |
| Pearson pred. gallery | Median rank | $571.7 \pm 50.0$ | $38.8 \pm 6.3$ | $144.2 \pm 27.8$ | $580.7 \pm 42.5$ |
| | Recall@10 (%) | $5.1 \pm 1.1$ | $22.3 \pm 2.4$ | $13.2 \pm 2.0$ | $7.0 \pm 0.8$ |

**Table S6 Paired RevPert versus Pearson ranks (Essential primary partition).** Per-query identity ranks for RevPert and Pearson on the same linear predicted gallery. One-sided Wilcoxon signed-rank alternative: Pearson rank > RevPert rank. Bootstrap 95% CI ($B$=2000) is on median(RevPert)−median(Pearson); negative favours RevPert. "RevPert better" counts queries with strictly lower RevPert rank.

| Line | Med. RevPert | Med. Pearson | $\Delta$ med. | 95% CI | $n$ | Wilcoxon $P$ | RevPert better |
|---|---|---|---|---|---|---|---|
| HepG2 | 266 | 572 | −306 | [−379.0, −235.0] | 593 | $1.9 \times 10^{-27}$ | 424/593 |
| K562 | 25 | 46 | −21 | [−30.0, −13.0] | 272 | $1.9 \times 10^{-19}$ | 190/272 |
| RPE1 | 90 | 185.5 | −95.5 | [−125.0, −64.5] | 384 | $1.6 \times 10^{-26}$ | 290/384 |
| Jurkat | 238.5 | 574.5 | −336 | [−393.0, −268.0] | 594 | $5.9 \times 10^{-30}$ | 419/594 |

**Table S7 Oracle and ESM-mean controls (Essential primary partition).** Observed-to-observed Pearson oracle is Supplementary-only (uses held-out $\Delta Y$ in the gallery). ESM-mean matching of top-magnitude genes to knockout sequence embeddings (UniProt sequences [4]; ESM2 [5]) is a weak prior and does not approach fair gallery methods.

| Line | Method | Median rank | Top-10 (%) | Top-1 (%) |
|---|---|---|---|---|
| HepG2 | RevPert | 266 | 12.5 | 0.84 |
| | Pearson pred. gallery | 572 | 3.7 | 0.2 |
| | ESM-mean topDEG→KO | 1206 | 0.7 | 0.2 |
| | Oracle obs. gallery | 1 | 100 | 100 |
| K562 | RevPert | 25 | 27.6 | 3.31 |
| | Pearson pred. gallery | 46 | 18.8 | 2.2 |
| | ESM-mean topDEG→KO | 504 | 1.1 | 0.7 |
| | Oracle obs. gallery | 1 | 100 | 100 |
| RPE1 | RevPert | 90 | 16.7 | 1.30 |
| | Pearson pred. gallery | 185.5 | 10.2 | 0.8 |
| | ESM-mean topDEG→KO | 789 | 1.3 | 0.3 |
| | Oracle obs. gallery | 1 | 100 | 100 |
| Jurkat | RevPert | 238.5 | 11.4 | 1.01 |
| | Pearson pred. gallery | 574.5 | 5.6 | 1.3 |
| | ESM-mean topDEG→KO | 1220 | 0.7 | 0.2 |
| | Oracle obs. gallery | 1 | 100 | 100 |

### S3 LINCS-KO expansions

LINCS-KO fairness audit, ten-line diagnostic means and per-line fold summaries for the main-text genetic identity comparisons.

**Table S8 LINCS-KO protocol fairness audit.** What is matched for the median-rank / Recall@10 comparison to official PDGrapher [2], and what is not. LINCS-KO screens derive from LINCS Level 3 expression [6].

| Dimension | Matched | Detail |
|---|---|---|
| Dataset | yes | LINCS-KO (LINCS L1000 genetic knockout) screens (10 lines); not Perturb-seq |
| Splits | yes | Official within-line random 5-fold genetic splits |
| Primary metrics | yes | Median rank of true intervention + Recall@10 |
| Query definition | yes | $\Delta Y$ = treated − diseased (post-knockout minus baseline) on the 10,716-gene PPI axis |
| Catalog restriction | yes | Genes restricted to those available to all compared methods |
| Gallery (matching / RevPert) | yes | Train-fold mean $\Delta Y$ by intervention gene |
| Official PDGrapher recipe | partial | Published genetic checkpoints scored under the identity-recovery prediction interface |
| Model class / objective | no | Residual gallery scorer versus GNN perturbation discovery |
| Pooled with Essential | no | Essential and LINCS-KO never pooled; primary baselines differ by dataset |

**Table S9 LINCS-KO diagnostics (ten-line means).** Mean median rank, Recall@10 (%), Recall@1 / partial% and nDCG across ten cell lines (five folds averaged per line, then mean over lines). Primary main-text genetic claims use median rank and Recall@10; this table reports PDGrapher-style secondary diagnostics without redefining the leaderboard.

| Method | Mean med. rank | R@10 (%) | R@1 / partial% | nDCG | $n$ lines |
|---|---|---|---|---|---|
| RevPert | **421.1** | **4.14** | **0.82** | **0.134** | 10 |
| CMap | 644.4 | 2.20 | 0.31 | 0.116 | 10 |
| GEM | 648.1 | 2.02 | 0.26 | 0.115 | 10 |
| Pearson | 691.8 | 2.18 | 0.31 | 0.115 | 10 |
| PDGrapher (official) | 4253.3 | 1.64 | 0.60 | 0.066 | 10 |

**Table S10 LINCS-KO per-line identity recovery (fold mean±s.d.).** Mean±s.d. of median rank (MR) and Recall@10 (%) across five official folds. Primary main-text genetic claims use the fold means; s.d. quantifies within-line fold variability.

| Cell | RevPert MR | RevPert R@10 | PDGrapher MR | PDGrapher R@10 | Pear. MR | Pear. R@10 |
|---|---|---|---|---|---|---|
| A375 | **361.8 ± 6.5** | 4.26 ± 0.36 | 5445.4 ± 113.9 | 0.75 ± 0.11 | 645.8 ± 17.5 | 2.56 ± 0.29 |
| A549 | **359.2 ± 9.1** | 4.86 ± 0.30 | 4354.2 ± 428.8 | 3.08 ± 0.41 | 576.0 ± 16.2 | 2.71 ± 0.14 |
| AGS | **411.6 ± 14.4** | 3.54 ± 0.31 | 3798.7 ± 140.9 | 1.33 ± 0.30 | 668.6 ± 10.6 | 2.09 ± 0.13 |
| BICR6 | **439.6 ± 23.2** | 3.59 ± 0.45 | 4583.6 ± 104.4 | 1.02 ± 0.27 | 737.5 ± 12.6 | 1.92 ± 0.25 |
| ES2 | **347.4 ± 15.8** | 6.38 ± 0.38 | 2769.9 ± 91.3 | 2.66 ± 0.26 | 615.0 ± 13.5 | 2.75 ± 0.21 |
| HT29 | **413.6 ± 25.8** | 3.66 ± 0.11 | 5623.2 ± 164.4 | 0.57 ± 0.07 | 631.6 ± 14.7 | 1.98 ± 0.06 |
| MCF7 | **494.3 ± 31.0** | 2.91 ± 0.26 | 3555.5 ± 215.3 | 1.75 ± 0.43 | 743.1 ± 24.4 | 1.70 ± 0.19 |
| PC3 | **558.3 ± 16.7** | 2.41 ± 0.15 | 3863.1 ± 83.7 | 1.31 ± 0.21 | 821.5 ± 28.0 | 1.34 ± 0.22 |
| U251MG | **345.0 ± 10.8** | 6.61 ± 0.30 | 3552.0 ± 936.0 | 3.33 ± 0.96 | 645.1 ± 13.2 | 2.98 ± 0.14 |
| YAPC | **480.4 ± 17.7** | 3.18 ± 0.23 | 4987.2 ± 114.8 | 0.63 ± 0.10 | 834.2 ± 16.3 | 1.73 ± 0.19 |

#### S4 Cross-context transfer and gallery diagnostics

Screen-external proving-ground analyses in the main text use lineage-matched galleries (HepG2 Essential for HCC; K562 GWPS for CML) [3]. To justify that design choice, we report a seed-1 diagnostic covering every Table S4 scorer. A model fitted in a *source* Essential line is applied to held-out identity queries from a *target* line. Expression-gallery scorers (Pearson, CMap [7], GEM [8], learn-only gallery-dual and fused RevPert) use the target linear predicted gallery [9] reindexed onto the source gene / PCA axis (missing genes set to zero). Ridge $\Delta Y \to P$ is fit on the source train split and ranks queries in the source $P$ atlas (queries restricted to knockouts present in that atlas). Diagonal entries are matched within-line controls and reproduce the Table S4 primary-partition medians; off-diagonal entries are transfers. Mean fused-RevPert median rank is 154.9 on matched diagonals versus 331.5 on the twelve transfer pairs (Recall@10 17.0% versus 9.5%). Learn-only encoding shows a similar drop (186.9 versus 396.4). Non-learned matchers change less on average because transfers into easy targets (especially K562) remain strong, but they do not replace matched within-line residual scoring on hard lines (HepG2 / Jurkat). Ridge transfers are unstable across $P$ atlases (cross-line mean median rank 610). A complementary forward-gallery identity board is Supplementary Table S12; the matching HCC dual-arm board is Supplementary Table S13.

**Table S11 Cross-context transfer of Table S4 scorers (Essential seed 1; median rank).** Source = training / gene-axis line; Target = query line. Lower is better. Matched controls (Source=Target) align with the main Essential leaderboard for Pearson / GEM / Ridge / learn-only / RevPert. CMap is the same mean-rank enrichment used in Figure 2c,d. Ridge uses the source $P$ atlas; other columns use the target linear predicted gallery on the source gene axis.

| Source | Target | Pearson | CMap | GEM | Ridge | Learn-only | RevPert |
|---|---|---|---|---|---|---|---|
| HepG2 | HepG2 | 572 | 668 | 587 | 638 | 289 | 266 |
| HepG2 | K562 | 47 | 54.5 | 53 | 999 | 138 | 44.5 |
| HepG2 | RPE1 | 167.5 | 221.5 | 162.5 | 1127 | 303.5 | 147.5 |
| HepG2 | Jurkat | 569 | 649 | 568 | 1026 | 581 | 451.5 |
| K562 | HepG2 | 576 | 668 | 587 | 96 | 683 | 610 |
| K562 | K562 | 46 | 53.5 | 48.5 | 45.5 | 35.5 | 25 |
| K562 | RPE1 | 169 | 232 | 163.5 | 260 | 238 | 176 |
| K562 | Jurkat | 577 | 638.5 | 565.5 | 285 | 583 | 559 |
| RPE1 | HepG2 | 657 | 605 | 650 | 143 | 671 | 658 |
| RPE1 | K562 | 46 | 63.5 | 45 | 422.5 | 117 | 45 |
| RPE1 | RPE1 | 185.5 | 215.5 | 195 | 180 | 169 | 90 |
| RPE1 | Jurkat | 598 | 639 | 593.5 | 361 | 510 | 560.5 |
| Jurkat | HepG2 | 610 | 695 | 618 | 460 | 586 | 540 |
| Jurkat | K562 | 48 | 53.5 | 51 | 1128.5 | 71.5 | 36 |
| Jurkat | RPE1 | 171 | 232 | 173.5 | 1017 | 274.5 | 149.5 |
| Jurkat | Jurkat | 574.5 | 653 | 565.5 | 708 | 254 | 238.5 |

**Table S12 Gallery-fidelity and residual-on-foreign-gallery checks (Essential seed 1).** Complements main-text Figures 2f and 4e: TxPert-reverse (Pearson) is the identity ceiling on held-out median rank, yet fails the HCC dual-arm anchors under the same Pearson scorer. Linear predicted RevPert is the primary seed-1 model (Table S4); remaining RevPert rows are residual models retrained on that gallery. Boldface marks the better scorer within a gallery when both are shown. scGPT coverage of the shared catalog is ∼38%; missing knockouts receive worst rank. Preferred-arm resistance ranks and gallery-collapse diagnostics follow in Supplementary Table S13.

| Forward gallery | Scorer | HepG2 | K562 | RPE1 | Jurkat |
|---|---|---|---|---|---|
| TxPert-GAT (within-line) | Pearson | **176** | **11** | **48** | **184** |
| | RevPert | 239 | 12 | 69.5 | 188.5 |
| Linear predicted | Pearson | 572 | 46 | 185.5 | 574.5 |
| | RevPert | **266** | **25** | **90** | **238.5** |
| TxPert x-cell LOO | Pearson | 792 | 170 | 447.5 | 868 |
| | RevPert | **600** | **79** | **366.5** | **671** |
| UniPert-ridge | Pearson | 991 | 219.5 | 486.5 | 945 |
| | RevPert | **850** | **184.5** | **474.5** | **779** |
| GEARS | Pearson | 1105 | **387.5** | 736 | 1031.5 |
| | RevPert | **1068** | 394.5 | **670.5** | **1000** |
| scGPT | Pearson | **1166** | **542.5** | **742.5** | **1172.5** |
| | RevPert | 1176 | 571.5 | 800 | 1175.5 |

The same foreign galleries are scored for HCC preferred-arm ranks under Pearson dual-arm matching (Supplementary Table S13); this is the numeric companion to main-text Figure 4e.

**Table S13 HCC preferred-arm dual-arm ranks across Essential forward galleries (seed 1).** Complements main-text Figure 4e. Scorer is Pearson dual-arm connectivity on each gallery; preferred arm is arg max of Arm A versus Arm B ranks (lower better). Expected literature arms: AURKA Arm B (GSE322742); METTL3 Arm A (GSE143233). † marks preferred-arm disagreement with that expectation. Gallery diagnostics: $n_{\text{unique}}$ distinct response spectra; frac_near0 fraction of near-zero-variance profiles. TxPert x-cell leave-one-line-out is included as a sensitivity row (not on the main identity–resistance board).

| Gallery | AURKA A | AURKA B | Pref. | METTL3 A | METTL3 B | Pref. | $n_{\text{unique}}$ | frac near0 |
|---|---|---|---|---|---|---|---|---|
| Linear predicted | 2369 | **4** | B | **6** | 2367 | A | 2373 | 0.000 |
| TxPert-GAT (within-line) | 150 | 638 | A† | 2099 | 275 | B† | 595 | 0.750 |
| TxPert x-cell LOO | 231 | 2143 | A† | 1482 | 892 | B† | 2373 | 0.000 |
| GEARS | 949 | 1425 | A† | **74** | 2300 | A | 2373 | 0.000 |
| UniPert-ridge | 487 | 1887 | A† | 1152 | 1222 | A | 2370 | 0.000 |
| scGPT | 842 | 61 | B | — | — | — | 902 | 0.000 |

### S5 Resistance proving-ground supplements

Signed nulls under Pearson geometry, CML dual-arm ranks and full RevPert versus Pearson / learn-only anchors for the main-text proving ground. Gallery-swap preferred-arm ranks that support Figure 4e are in Supplementary Note S4 (Supplementary Table S13).

**Table S14 Proving-ground signed nulls and literature-anchor checks (Pearson geometry).** HCC: HepG2 Essential gallery [3]; CML: K562 GWPS ($n = 9866$) [3]. All ranks and permutation $P$ values in this table are Pearson dual-arm scores, not fused RevPert. Headline fused ranks are in main-text Table 1 and Supplementary Table S16 (AURKA 2331/29; METTL3 65 on Arm A; CML on-drug 1/16/18/27/34). Near-zero Top-50 overlaps mean the Pearson reverse shortlist is not just the biggest DEGs, and that direction matters. CML Top-50 nulls across all three IR contrasts are shown in Supplementary Fig. S1b; $P$ values are one-sided preferred-arm ranks under 500 gene-axis permutations of $\Delta Y^\star$ (null median preferred rank ∼2500).

| Signature | What was tested | Value |
|---|---|---|
| GSE322742 HepG2×sorafenib | Arm A Top-50 overlap with top \|DEG\| genes | 0 |
| GSE322742 HepG2×sorafenib | Arm A Top-50 after flipping $\Delta Y^\star$ | 0 |
| GSE322742 HepG2×sorafenib | AURKA Pearson ranks (Arm A / Arm B) | 2370 / **4** |
| GSE143233 patient SR vs normal | METTL3 Pearson ranks (Arm A / Arm B) | **6** / 2368 |
| GSE143233 patient SR vs normal | Either-arm Top-50 overlap with top \|DEG\| genes | 0 |
| GSE120932 IR + drug | MYB Pearson preferred rank / permutation $P$ | **1** / $< 0.002$ |
| GSE120932 IR + drug | STAT5A Pearson preferred rank / permutation $P$ | 13 / $< 0.002$ |
| GSE120932 IR + drug | RUNX1 Pearson preferred rank / permutation $P$ | 20 / 0.006 |
| GSE120932 IR + drug | STAT5B Pearson preferred rank / permutation $P$ | 25 / 0.002 |
| GSE120932 IR + drug | BCR Pearson preferred rank / permutation $P$ | 36 / 0.006 |
| GSE120932 IR + drug | ABL1 Pearson preferred rank / permutation $P$ (mid-catalog control) | 3206 / 0.62 |

**Table S15 CML dual-arm ranks on K562 GWPS ($n = 9866$) [3].** Pearson Arm A / Arm B ranks for pre-specified CML pathway anchors [10] on the three GSE120932 contrasts [11], complementary to Fig. 3b. Arm A is phenocopy of the query; Arm B is the opposite signed match. The two ranks of an anchor sum to $n+1$ by construction; boldface marks the better arm. ABL1 stays mid-catalog on all three contrasts (range over contrasts). These genes are largely absent from the smaller K562 Essential gallery. Fused RevPert preferred-arm ranks are in Supplementary Table S16.

| Signature | Anchor | Arm A | Arm B |
|---|---|---|---|
| GSE120932 IR + drug | MYB | **1** | 9866 |
| GSE120932 IR + drug | STAT5A | 9854 | **13** |
| GSE120932 IR + drug | RUNX1 | 9847 | **20** |
| GSE120932 IR + drug | STAT5B | **25** | 9842 |
| GSE120932 IR + drug | BCR | **36** | 9831 |
| GSE120932 IR off drug | STAT5B | 29 | 9838 |
| GSE120932 IR off drug | STAT5A | 9823 | 44 |
| GSE120932 IR off drug | MYB | 96 | 9771 |
| GSE120932 IR off drug | BCR | 179 | 9688 |
| GSE120932 IR off drug | RUNX1 | 9374 | 493 |
| GSE120932 spindle IR | MYB | **1** | 9866 |
| GSE120932 spindle IR | STAT5A | 9829 | 38 |
| GSE120932 spindle IR | RUNX1 | 9819 | 48 |
| GSE120932 spindle IR | STAT5B | 87 | 9780 |
| GSE120932 spindle IR | BCR | 108 | 9759 |
| GSE120932 (all three) | ABL1 | 2577–3501 | 6366–7290 |

**Table S16 Full RevPert on HCC and CML proving-ground anchors.** Preferred-arm ranks under Pearson, learn-only and fused RevPert. HCC uses HepG2 Essential residual training (seed 1) [3,12,13]; CML uses a GWPS gallery-native residual model scored on all three GSE120932 IR contrasts [3,11]. Learn-only displaces anchors; fused RevPert stays near Pearson.

| Contrast | Anchor | Pearson | Learn-only | RevPert | Pref. arm (RevPert) |
|---|---|---|---|---|---|
| GSE322742 HepG2 SR−parental | AURKA | 4 | 218 | 29 | B |
| GSE143233 patient SR−normal | METTL3 | 6 | 2127 | 65 | A |
| GSE120932 IR + drug | MYB | 1 | 3 | 1 | A |
| GSE120932 IR + drug | STAT5A | 13 | 921 | 16 | B |
| GSE120932 IR + drug | RUNX1 | 20 | 116 | 18 | B |
| GSE120932 IR + drug | STAT5B | 25 | 198 | 27 | A |
| GSE120932 IR + drug | BCR | 36 | 140 | 34 | A |
| GSE120932 IR off drug | MYB | 96 | 5 | 64 | A |
| GSE120932 IR off drug | STAT5B | 29 | 37 | 21 | A |
| GSE120932 IR off drug | STAT5A | 44 | 664 | 47 | B |
| GSE120932 spindle IR | MYB | 1 | 7 | 1 | A |
| GSE120932 spindle IR | STAT5A | 38 | 1132 | 47 | B |
| GSE120932 spindle IR | RUNX1 | 48 | 178 | 43 | B |
| GSE120932 spindle IR | BCR | 108 | 230 | 96 | A |

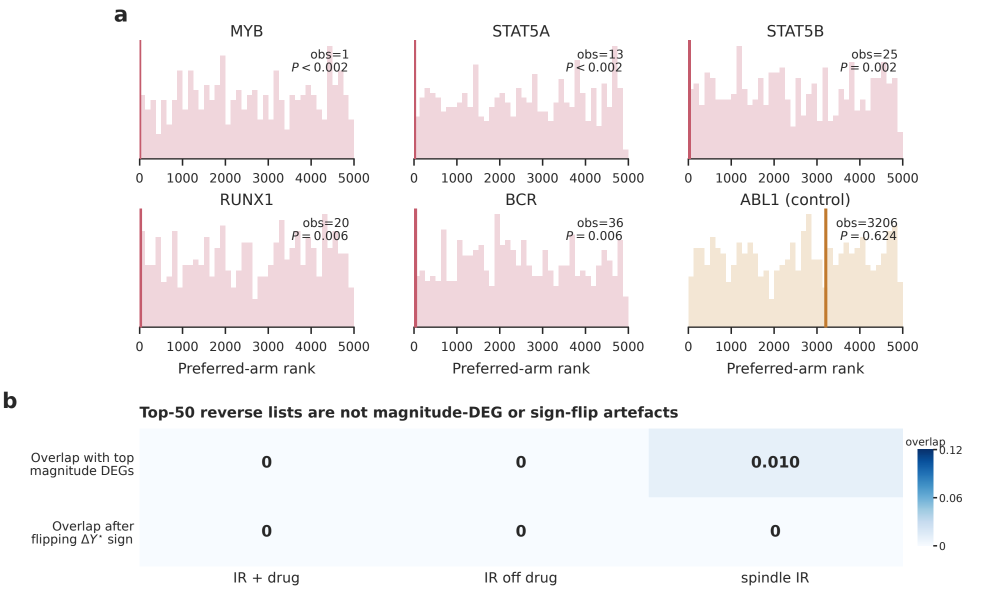


**Fig. S1 CML signed nulls on K562 GWPS under Pearson geometry (SI-only).** **a**, Pearson preferred-arm rank under the on-drug GSE120932 contrast versus 500 gene-axis permutations of $\Delta Y^\star$ (histograms); vertical lines mark observed Pearson ranks for five pathway anchors and mid-catalog ABL1. These nulls do not test the fused GWPS residual (on-drug RevPert ranks 1/16/18/27/34; Supplementary Table S16). **b**, Across the three IR contrasts, Pearson Arm A Top-50 lists barely overlap top-magnitude DEGs or the Arm A list after sign-flipping $\Delta Y^\star$ (near 0 passes). Numeric summaries are in Supplementary Table S14.

### S6 Model hyperparameters

Default training and architecture settings for Essential, LINCS-KO and the GWPS proving-ground residual model.

**Table S17 RevPert training and architecture defaults.** Essential models were trained independently for each cell line and partition; LINCS-KO models were trained independently within each official fold.

| Component | Setting |
|---|---|
| Profile PCA dimension | 256 (fit on training profiles only) |
| Shared profile MLP | 512–512 hidden; 128-d $\ell_2$-normalised output; GELU; dropout 0.1 |
| Fused score | $z(\text{Pearson}) + \text{softplus}(\beta)\, z(\text{learned similarity})$ |
| Residual initialization | $\beta = -1$ ($\alpha \approx 0.31$) |
| Loss | Full-gallery fused InfoNCE [14] + Pearson-teacher regularizer (weight 0.25) |
| Optimiser | AdamW ($10^{-3}$, weight decay $10^{-4}$) |
| Schedule | Cosine learning-rate decay |
| Essential batch / epochs | 64 / 40 |
| PDGrapher batch / epochs | 256 / 25 |
| Checkpoint select | Validation fused MRR with learned-score tie-break (Essential and LINCS-KO) |
| Selected $\alpha^\star$ (HepG2, seed 1) | 0.44 |
| GWPS transductive model | Full observed catalog as gallery; noise-augmented validation saturates (median rank 1); deployed $\alpha = 0.1$ from leave-self-out retune; no held-out identity validation |
| Ridge baseline $\alpha$ | 1 |
